# Hydrocephalus caused by *Katnip* deletion is linked to increased ciliogenesis and reduced proliferation of neuroprogenitor cells

**DOI:** 10.64898/2026.05.01.722314

**Authors:** Ana Limerick, Cheuk Ying Chu, Jacob S. Turner, David L. Brautigan, Wenhao Xu, Zheng Fu

**Affiliations:** Department of Pharmacology, University of Virginia School of Medicine, Charlottesville, Virginia; Department of Microbiology, Immunology, and Cancer Biology, University of Virginia School of Medicine, Charlottesville, Virginia; Genetically Engineered Murine Model Core, University of Virginia School of Medicine, Charlottesville, Virginia

**Keywords:** primary cilia, neuroprogenitor cells, brain development, ciliopathy, ciliogenesis

## Abstract

**Background:** *KATNIP* (Katanin-interacting protein), also known as *KIAA0556*, is one of the human genes with pathogenic variants linked to Joubert syndrome, an archetypal neurodevelopmental ciliopathy. KATNIP is a scaffolding protein with a critical role in ciliogenesis. In this study, we characterized the ciliopathy phenotypes due to *KATNIP* gene deletion.

**Results:** We produced a *Katnip* null mouse model using CRISPR-Cas12a (Cpf1). The null heterozygotes appeared normal while the homozygotes died around postnatal day 9, showing severe hydrocephalus and deficiency in neuroprogenitor cell proliferation. Katnip-deficient cells in the brain have a higher rate of cilia formation and longer cilia than wild type cells.

**Conclusion:** KATNIP loss of function gives rise to hydrocephalus found in Joubert syndrome. The results indicate that KATNIP restricts ciliogenesis and cilia extension and supports proliferation of neuroprogenitor cells in the brain.

## 1. INTRODUCTION

Joubert syndrome (JS) is an inherited ciliopathy characterized by abnormal brain development and is associated with mutations in more than 30 genes encoding proteins with a role in the structure and function of the primary cilium.^1,2^ Most mammalian cells possess a primary cilium, a dynamic microtubule-based apical membrane protrusion originated from centrioles.^3^ The cilium enables cells to sense environmental cues and integrate signaling responses through compartmentalization and support of receptor tyrosine kinases, G-protein coupled receptors, and second messengers.^4^ The primary cilium also plays a critical role in cell cycle as a structural checkpoint.^5^ Before mitosis, cilia resorption is required to release centrosomal components essential for spindle formation. Following mitosis, primary cilia are reassembled to resume their sensory and signaling functions during interphase. This dynamic cycle of resorption and reformation involves regulated microtubule reorganization, which enables ciliary length changes through microtubule disassembly and reassembly.^6^ Disruption of this process may impede cell cycle, mitotic entry and cell replication.^7^

Mutations in *KATNIP*, a human gene encoding katanin-interacting protein, have been identified in individuals with Joubert syndrome.^8^ These include four frameshift and truncation variants: N74fs*, Q892*, R1253Qfs*, and M1474Cfs*.^9-12^ The only recognizable features in KATNIP protein are three domains of unknown function (DUFs), as annotated by the NCBI Conserved Domain Database.^10,13^ Each of the identified mutations introduces a premature stop codon, resulting in truncated proteins.^14^ KATNIP functions as a scaffold protein at the cilium base (the basal body) interacting with multiple protein partners.^10,15^ Recently, we published in vitro evidence supporting a critical role of KATNIP in the control of cilia formation and elongation through stabilization and activation of CILK1 (ciliogenesis-associated kinase 1).^13,14^ However, the function of KATNIP in mammalian development remains poorly understood. Specifically, it is unclear how KATNIP contributes to primary cilium regulation, neuroprogenitor proliferation, and brain development. Prior studies have not established a causal link between KATNIP dysfunction and the neurodevelopmental phenotype observed in JS patients and the cellular mechanisms underlying this link.

To address this gap in our knowledge, we generated a *Katnip* knockout (KO) mouse model to mimic the KATNIP deficiency in Joubert syndrome. This study aimed to characterize the developmental and cellular phenotypes associated with the loss of KATNIP and to examine the role of KATNIP in coordinating cilia dynamics and cell cycle progression. Our results herein demonstrate that *Katnip* deletion leads to severe hydrocephalus. Hydrocephalus is only seen in a distinct subtype of JS patients.^16,17^ The genetic basis of this association remains elusive. Although a truncating mutation in *KATNIP* was reported as one of two independent causative mutations in a family of JS patients with hydrocephalus, it has yet to be determined if this truncated KATNIP variant is the primary cause of hydrocephalus.^11^

Hydrocephalus, characterized by abnormal accumulation of CSF in the brain ventricles, is a structural defect in the brain commonly found in ciliopathies.^18^ The conventional “Standard Model of Hydrocephalus” attributes this phenotype to defective motile cilia in ependymal cells, which are proposed to impair CSF circulation and lead to ventricular enlargement.^19^ However, clinical evidence suggests that motile cilia dysfunction plays a minimal role in human hydrocephalus, and motile ciliopathies rarely result in this condition. A revised model proposes that hydrocephalus arises from intrinsic defects in brain development, rather than disrupted CSF flow.^20^ Specifically, genetic mutations that impair neural stem cell proliferation and cortical neurogenesis result in a hypoplastic cortex with reduced tissue elasticity and stiffness. This mechanically unstable cortex cannot resist the pressure of CSF, leading to ventricular expansion. Supporting this model, mouse models with defective primary (non-motile) cilia exhibit hydrocephalus, caused by impaired neuroprogenitor proliferation and neurogenesis. Our results herein reinforced this model and linked *KATNIP* variants to the hydrocephalus phenotype of Joubert syndrome.

## 2. RESULTS

### 2.1 Katnip deficiency causes severe hydrocephalus and perinatal lethality

We used CRISPR/Cas12a to genetically engineer deletion mutations in exon 4 of mouse *Katnip* (Fig. 1A). Sequencing data confirmed a 25-base pair deletion around exon 4 of the *Katnip* gene, resulting in a frameshift and a premature stop codon. This truncated, short transcript is likely unstable and degraded by cellular surveillance systems. Genotyping by PCR confirmed the deletion, with the mutant amplicon ∼25 bp shorter than the wild type (WT) product (Figure 1B). Western blot analysis further demonstrated the absence of full-length Katnip protein (∼200 kDa) in brain tissue lysates from *Katnip* homozygous KO mice, confirming successful ablation of Katnip (Figure 1C).

**Figure 1:**
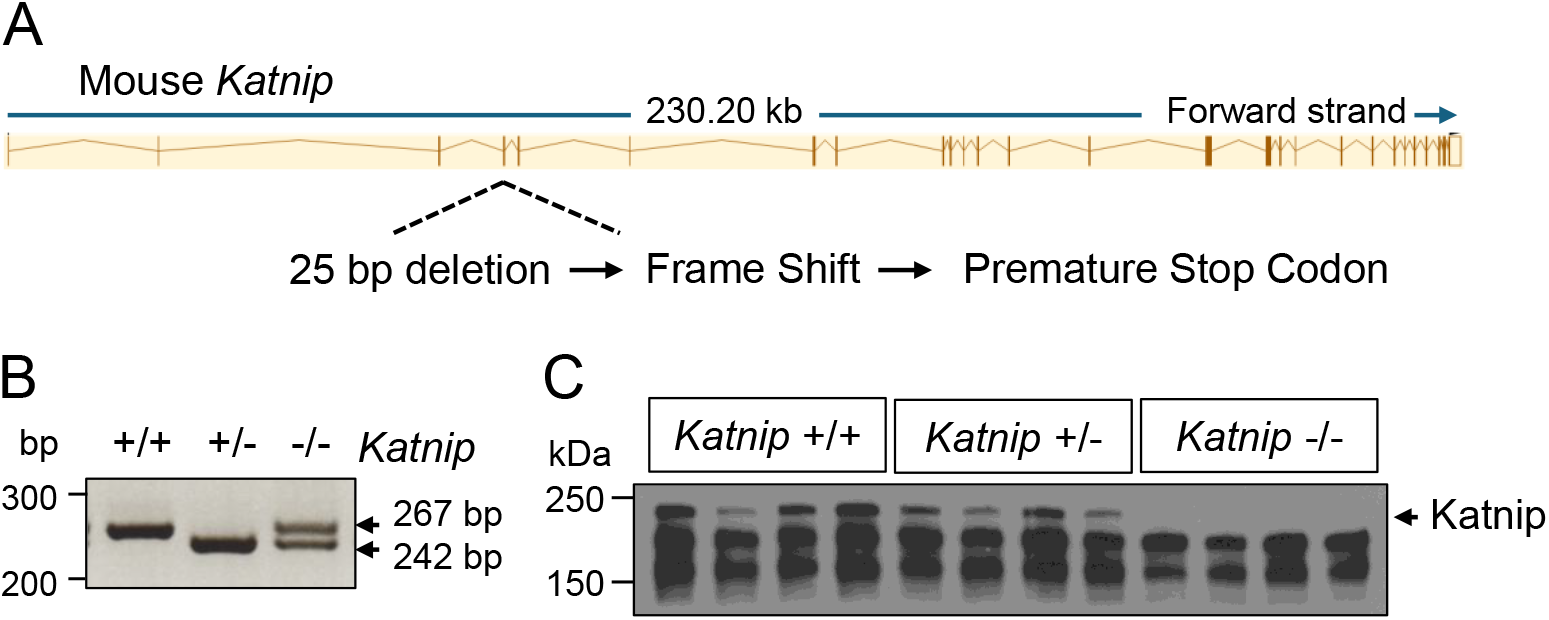
Generation of a *Katnip* knockout mouse model. (A) Schematic illustration of CRISPR/Cas12a-mediated knockout of *Katnip*. (B) Genotyping PCR results showing distinct patterns of the wild type (WT, 267 bp) and the knockout (KO, 242 bp) bands between three different genotypes. (C) A Western blot analysis of Katnip protein in mouse brain tissues from three different genotypes.

The founder mice carrying *Katnip* heterozygous deletion were viable and fertile, and indistinguishable from their wild type littermates in appearance and behavior. This result can be explained by the lack of noticeable differences in the abundance of Katnip protein between the heterozygotes and the wild types (Fig. 1C). By contrast, the null homozygotes displayed dramatic physical deformities. Compared with WT littermates, they were significantly smaller in size, unsteady in their movements (Fig. S1), and failed to survive past postnatal day 10. We focused our subsequent phenotypic analysis on the homozygotes, hereinafter referred to as KO mice. At postnatal day 9, KO mice exhibited markedly reduced body size compared to WT littermates (Fig. 2A), with a 40% reduction in body length (P < 0.001, Fig. 2B). Upon brain dissection, we observed a prominent hydrocephalus phenotype characterized by a substantial buildup of cerebrospinal fluid (CSF) in the brain ventricles. Histology corroborated this observation, showing extensive dilation of the lateral ventricles in KO brains relative to WT controls (Fig. 2C). These results demonstrate that loss of Katnip results in perinatal lethality, severe physical disabilities (e.g. loss of control in balance and coordination in walking) and structural defects in the brain (e.g. gross enlargement of lateral ventricles).

**Figure 2:**
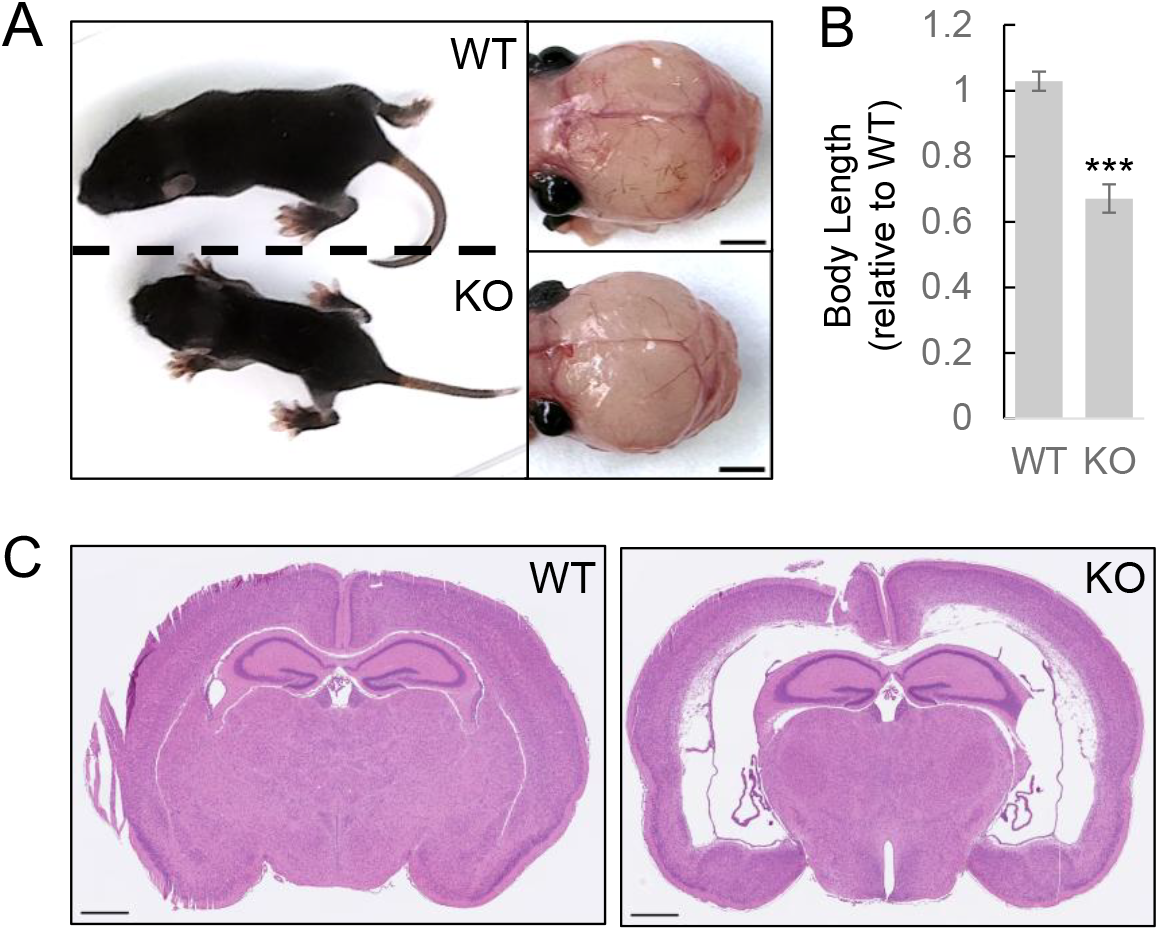
Homozygous depletion of *Katnip* produces a perinatal lethal hydrocephalic phenotype. (A) Representative images of the whole body and the dorsal view of dissected brains from the WT control and the homozygous null littermate at postnatal day 9 (P9). (B) Shown are relative body length, *n* = 4 mice, P9, average ± SD, two-sided unpaired Student’s *t* test, ***P < 0.001. (C) Hematoxylin and eosin (H&E) staining of coronal mouse brain sections, showing broad expansion of the lateral ventricles in the homozygote brain. Scale bar, 500 µm.

### 2.2 Loss of Katnip impedes cell cycle and neuroprogenitor cell proliferation

We investigated if the hydrocephalus phenotype of Katnip KO is due to impaired brain development associated with primary cilia dysfunction. Given that KATNIP suppresses cilia formation *in vitro*, we hypothesized that loss of KATNIP *in vivo* would promote ciliogenesis. This would enhance the ciliary cell cycle checkpoint, thus hindering cell cycle progression and replication of neuroprogenitor cells during brain development. To test this hypothesis, we examined proliferation of WT and Katnip-deficient neuroprogenitor cells in the hippocampus. We chose hippocampus to avoid the major structural disruptions in the cerebral cortex. We immunostained cells in the dentate gyrus of the hippocampus of postnatal day 9 (P9) brains (Fig. 3A). Sox2 was used to identify neuronal progenitor cells, while phospho-histone H3 (pH3) and Ki67 were markers for mitotic and proliferating cells, respectively. We quantified the proportion of Sox2+ cells that were also positive for pH3 or Ki67 to determine the mitotic index and overall proliferation rate of the neuronal progenitor cell population in the hippocampus.

**Figure 3:**
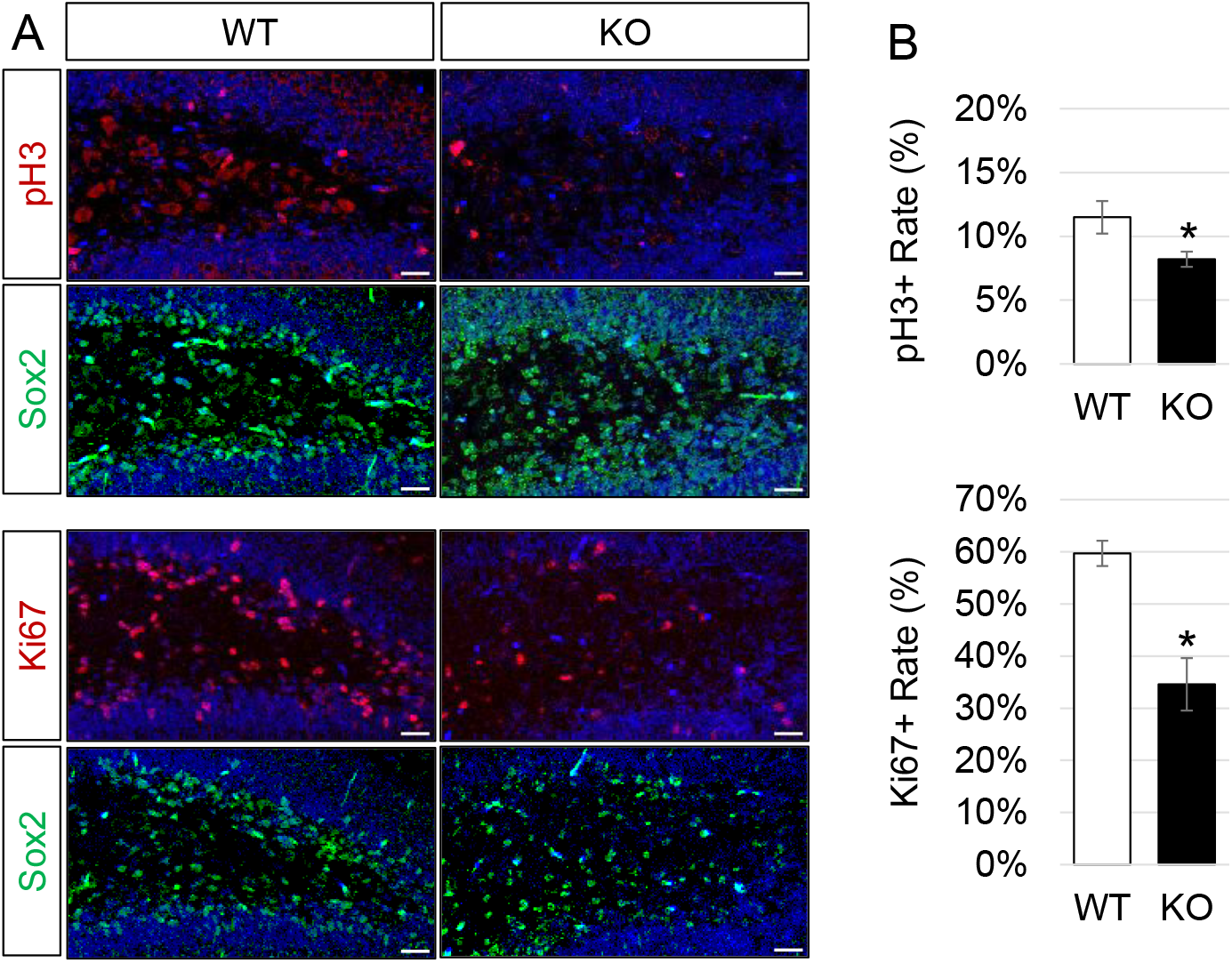
Katnip deficiency decreases the proliferation of neuroprogenitor cells. (A) Representative fluorescence microscopy images showing phospho-histone H3 (pH3)-positive mitotic cells (red), Ki67-positive proliferating cells (red), and Sox2-positive neuroprogenitor cells (green) in the dentate gyrus of wild type (WT) and homozygous knockout (KO) mice. DAPI stains nuclei (blue). Scale bar, 5 µm. (B) Quantification of pH3+ and Ki67+ rates in Sox2+ neuroprogenitor cells from WT and homozygous KO brains (*n* = 3 mice, P9). Shown are average ± SD, two-sided unpaired Student’s *t* test, *P < 0.05.

Quantification in Figure 3B demonstrates a statistically significant reduction in both pH3+/Sox2+ and Ki67+/Sox2+ neuronal progenitor cell populations in *Katnip* KO brains compared to WT brains. Specifically, the proportion of Sox2+ neuronal progenitors that were also pH3+, marking mitotic cells, decreased from 11 ± 2% in WT to 8 ± 1% in KO (p < 0.05), a ∼27% reduction. Similarly, the proportion of Ki67+/Sox2+ cells, reflecting the overall proliferation of neuroprogenitor cells, declined from 60 ± 3% in WT to 35 ± 5% in KO (p < 0.01), a ∼42% reduction.

Overall, *Katnip* deletion significantly reduced the proliferation and fraction of mitotic neuroprogenitor cells, consistent with the revised model of hydrocephalus, characterized by impaired neuroprogenitor proliferation and neurogenesis.

### 2.3 Katnip deficiency increases ciliation rate and cilia length

Dynamic regulation of cilia length and cilia disassembly is essential for neuronal progenitor proliferation. Because cells must resorb cilia before entering mitosis, an increase in ciliation rate or cilia length may hinder cell cycle progression and contribute to reduced neurogenesis. To assess whether Katnip regulates cilia dynamics, we examined cilia structure in the dentate gyrus of the hippocampus of P9 brains using immunofluorescence.

As shown in Figure 4A, we stained for Arl13b (green) to label the ciliary axoneme, FOP (red) to mark centrioles, and DAPI (blue) for nuclei. Quantification across three littermate pairs revealed a statistically significant increase in both the percentage of ciliated cells and the average cilia length in *Katnip* KO cells compared to WT (Fig. 4B). The ciliation rate increased from 39 ± 2% in WT to 65 ± 7% in KO (p < 0.01), reflecting a ∼67% increase in the proportion of cells bearing primary cilia. Likewise, the average cilia length rose significantly from 1.13 μm in WT to 1.64 μm in KO (p < 0.01), a ∼45% increase. We conclude that loss of *Katnip* expression promotes the prevalence and elongation of primary cilia, supporting the notion that impaired ciliary disassembly contributes to enhanced ciliary checkpoint and reduced cell proliferation.

**Figure 4:**
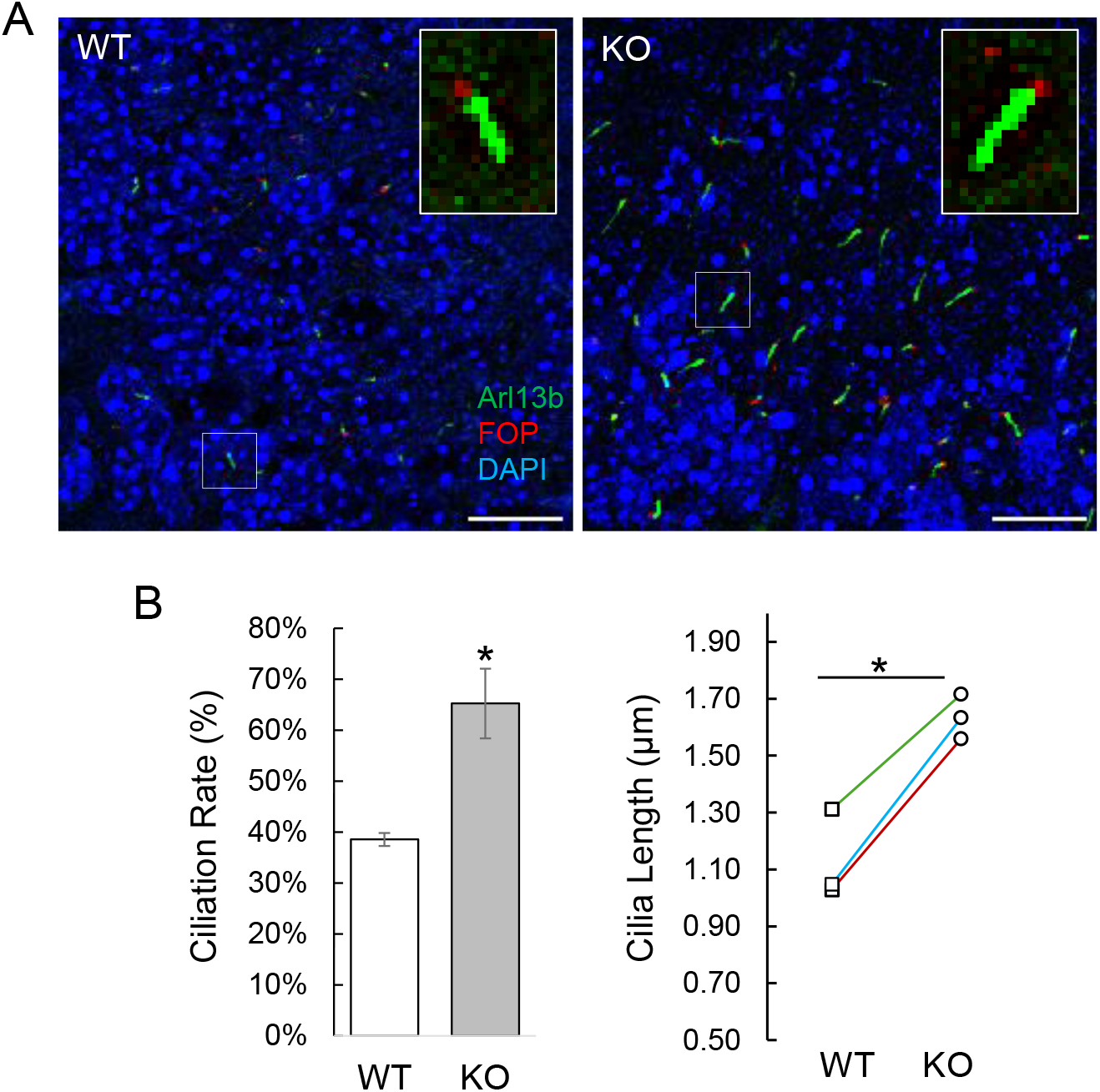
Loss of Katnip increases primary cilia frequency and length in the brain. (A) Representative images of primary cilia from WT control and KO mutant mouse hippocampal sections. Cilia axoneme stained with Arl13B (green), basal body stained with FOP (red), and nuclei stained with DAPI (blue). Scale bar, 5 µm. (B). Quantification of cilia frequency and cilia length in hippocampal tissue sections between three pairs of KO mutants and WT littermates (*n* = 3 mice, P9). Shown are average ± SD, two-sided Student’s *t* test, *P<0.05.

## 3. DISCUSSION

*KATNIP* variants have been found in patients with Joubert syndrome and our goal in this study was to determine whether disruption of KATNIP functions result in the characteristic phenotypes of this archetypal neurodevelopmental ciliopathy and whether compromised ciliogenesis and proliferation of neuroprogenitor cells contribute to the ciliopathy phenotypes. Using a CRISPR/Cas12a-engineered mouse model with two *Katnip* null alleles, we demonstrated that *Katnip* deficiency causes severe hydrocephalus and physical disabilities, leading to perinatal lethality. The results further revealed that loss of Katnip increased primary cilia frequency and length and decreased the mitotic rate and proliferation of neuroprogenitor cells. These findings together support a model in which KATNIP plays a pivotal role in the regulation of primary cilia and cell cycle of neuroprogenitor cells during neurogenesis. Loss of KATNIP function results in phenotypes that resemble Joubert syndrome.

Brain phenotypes were only detected in mice carrying both *Katnip* null alleles, indicating that a null mutation in *Katnip* is recessive. *KATNIP* truncation mutations are also recessive in producing ciliopathy phenotypes in patients with Joubert syndrome. These results together suggest that one functional allele of *KATNIP* is sufficient to support primary cilia dynamics and neuroprogenitor cell proliferation.

This work expands our understanding of the cellular mechanisms underlying KATNIP dysfunctions in Joubert syndrome. While previous studies identified *KATNIP* mutations in affected individuals, the biological role of this gene in neurodevelopment had not been experimentally validated. Our findings provide direct in vivo evidence that KATNIP is required for the maintenance of primary cilia, a key structural checkpoint in cell cycle progression into mitosis for neural stem/progenitor cells. This mechanistic insight links impaired cilia dynamics and cell cycle to defective neurogenesis and hydrocephalus, supporting a revised model in which primary cilia dysfunction and abnormal neurogenesis underlies ventricular enlargement.

In a previous study, a mutant mouse with the genetrap integrated after exon 14 of *Katnip* also exhibited hydrocephalus with ventricular distension.^10^ Ependymal motile cilia in lateral ventricles of mutant mice seemed normal in their ability to beat and move cerebral spinal fluid as compared with wild type mice. The authors concluded that the pathogenic mechanism underlying KATNIP-associated hydrocephalus may not involve ependymal motile cilia. Our results here suggest that abnormal development of neuroprogenitor cells may underlie neonatal hydrocephalus phenotype caused by KATNIP deficiency, similar to what was reported in a mouse models of Bardet-Biedl syndrome.^21^

Our analyses of cilia and developmental phenotypes associated with *Katnip* null mutations were not exhaustive and have several limitations. First, due to massive structural disruption in the cortical subventricular zone, our analysis focused on the neuroprogenitor cell population in dentate gyrus of the hippocampus. Knowledge about the molecular and cellular events leading to a hypoplastic cortex must be gained through more in-depth analyses of Katnip-deficient brains earlier during embryonic development. Second, we could not provide direct evidence for reorganization of axonemal MTs in *Katnip* KO cells because the size of the cilia axoneme and density of MTs pose significant technical hurdles for directly visualizing MT severing events in situ. It is worth mentioning that we have preliminary evidence to indicate that the number of EB-1 labelled MT plus ends is markedly reduced in the cytoplasm of *Katnip* null cells (data not shown), which suggests a significant alteration of MT dynamics in the absence of KATNIP. Third, we have not determined how *Katnip* deletion impacts organ systems other than brain. Although ciliopathies affect multi-organ systems, Joubert syndrome is primarily a neurodevelopmental disorder. In this study, we focused our analyses on the brain that was profoundly affected by the loss of Katnip. However, we cannot exclude the possibility that impact on other organ systems may also contribute to the perinatal lethality of Katnip KO. Fourth, we are fully aware that alterations of primary cilia number and length can affect not only cell cycle progression but also cilia-dependent signal transduction, such as the Hedgehog signaling. It is possible that combined effects of altered cell cycle and ciliary Hedgehog signaling lead to tissue structural defects in Katnip null brains. Fifth, this study did not examine whether disruption of KATNIP functions outside the primary cilium contributes to the neurodevelopmental defects. A recent study from *Dictyostelium* revealed an extraciliary role of KATNIP in the maintenance of MT-based endosomal trafficking. The detailed mechanisms underlying the hydrocephalic phenotype and the perinatal lethality merits further investigation.

Our current working model proposes that KATNIP functions as a scaffolding protein that stabilizes CILK1, a ciliary kinase, which in turn phosphorylates katanin, a microtubule (MT)-severing enzyme. Since axonemal microtubules provide the structural support to the primary cilium and MT reorganization is the molecular basis of cilia dynamics, this KATNIP-CILK1-katanin axis may provide a direct mechanism by which KATNIP regulates MT-based cilia assembly and disassembly during cell cycle. Proteomic analyses suggest complexes of these proteins, but direct *in vivo* functional validation remains incomplete. Future experiments will be critical to determine whether KATNIP is required for CILK1-mediated phosphorylation of katanin and whether this pathway is necessary for Katanin-mediated MT severing, cilia disassembly, and cell cycle progression during brain development.

## 4. EXPERIMENTAL PROCEDURES

### 4.1 Katnip knockout mouse model

All procedures involving animals were performed in accordance with ethical standards in an animal protocol that was approved by the Institutional Animal Care and Use Committee. The CRISPR-Cas12a technology was used to generate *Katnip* knockout founder mice. A sgRNA (TCTTCCAGGATACTAAAGCATTT) is selected based on the search via the CRISPR guide design algorithm CRISPOR (http://crispor.tefor.net/). The deletion mutations were introduced into the exon 4 of the wild-type *Katnip* allele. Reagents crRNA and Cas12a were purchased from IDT (Coralville, IA). Ribonucleic protein (RNP) complex was formed by mixing and incubating Cas12a at 0.5 μg/μl with crRNA at 0.15 μg/μl in RNase-free microinjection buffer (10 mM of Tris-HCl, pH 7.4, 0.25 mM of EDTA) at 37°C for 10 minutes. Fertilized eggs were collected from B6SJLF1 females mated with males. RNP were delivered into fertilized eggs by electroporation with a NEPA21 super electroporator (Nepa Gene Co., Ltd. Chiba, Japan) under the following conditions: 2 pulses at 40 V for 3 msec with 50 msec interval for poring phase; 3 pulses at 7 V for 50 msec with 50 msec interval for transferring phase. The zapped zygotes were cultured overnight in KSOM medium (EMD Millipore, Billerica, MA) at 37°C in 5% CO2. The next morning, zygotes that had reached the two-cell stage were implanted into the oviducts of pseudo pregnant foster mothers of ICR strain (Envigo, Indianapolis, IN). Pups born to the foster mothers were screened using tail snip DNA by PCR genotyping and restriction digestion analysis followed by Sanger’s sequencing. Germline transmission of the desired alleles was confirmed by breeding the founders with wild type C57BL/6J mice. Animals were housed in a temperature-controlled colony room on a 12-hour light cycle and had access to food and water ad libitum. Aging breeder mice were euthanized by CO_2_ inhalation. Experimental mice were euthanized by cervical dislocation before tissue harvest for histological and morphometric analysis.

### 4.2 Western blot

Brains were extracted from both male and female mice (postnatal day 9) and snap-frozen in liquid nitrogen. Brain tissues were homogenized in ice-cold lysis buffer (50 mM Tris-HCl, pH 7.4, 150 mM NaCl, 1% NP-40, 2 mM EGTA, and complete protease and phosphatase inhibitors from Roche, Basel, Switzerland). Tissue homogenates were cleared by centrifugation, boiled for 5 min in an equal volume of 2X Laemmli sample buffer (120 mM Tris-HCl, pH 6.8, 4% SDS, 20% glycerol, 10% β-mercaptoethanol, 0.02% bromophenol blue), and loaded on an SDS-polyacrylamide gel. Following electrophoresis protein samples were transferred to a PVDF membrane and blocked for one hour in 5% dry milk before incubation with primary antibodies in TBS containing 0.1% Tween-20 and 5% bovine serum albumin (BSA) for overnight at 4°C. This was followed by multiple rinses and one-hour incubation with horseradish peroxidase (HRP)-conjugated secondary antibody. Chemiluminescent signals were developed using Millipore Immobilon ECL reagents (EMD Millipore) according to manufacturer’s instructions. The KATNIP rabbit polyclonal antibody was generated against the human KATNIP peptide x at Genscript (Piscataway, NJ, USA).

### 4.3 Histology

Whole brains were dissected from mice of both sexes (postnatal day 9). Isolated brains were fixed in 10% neutral buffer formalin overnight, rinsed in PBS, and then paraffin embedded at the Research Histology Core of University of Virginia. Coronal sections were stained with H&E (Hematoxylin and Eosin) for histological analysis.

### 4.4 Confocal immunofluorescence microcopy

After heat-induced antigen retrieval in Tris-EDTA (pH 8.0), brain tissue sections were rinsed in PBS, and then permeabilized by 0.2% Triton X-100 in PBS. After one hour in blocking buffer (3% goat serum, 0.2% Triton X-100 in PBS), tissues were incubated in primary antibody at 4°C overnight followed by rinses in PBS and one hour incubation with Alexa Fluor-conjugated goat anti-rabbit or anti-mouse IgG secondary antibody (Abcam, Cambridge, MA, ab150084 and ab150115). After multiple rinses, slides were mounted in antifade reagent containing DAPI (4’, 6-diamidino-2-phenylindole) for imaging via ZEISS LSM 700 Confocal Microscope at the UVA Advanced Microscopy Facility.

The following primary antibodies were used in this study: cilia marker Arl13B mouse monoclonal antibody (ProteinTech, Rosemont, IL, 17711-1-AP), FOP rabbit polyclonal antibody (GeneTex, Irvine, CA, GTX46056), Sox2 mouse monoclonal antibody (ProteinTech, Rosemont, IL, 66200-1), Phospho-histone H3 rabbit antibody (Cell Signaling Technology, Danvers, MA, #8507), Ki67 rabbit polyclonal (ProteinTech, Rosemont, IL, 13967-1-AP).

### 4.5 Cilia length measurement

Zen 2009 program was used with the confocal Laser Scanning Microscope 700 (LSM 700) to collect z stacks at 0.5 μm intervals to incorporate the full axoneme based on cilia marker Arl13b staining. All cilia were then measured in Fiji/ImageJ (version 1.52t) via a standardized method based on the Pythagorean Theorem in which cilia length was based on the equation L2 = z2 + c2, in which “c” is the longest flat length measured of the z slices and “z” is the number of z slices in which the measured cilia was present multiplied by the z stack interval (0.5 μm).

### 4.6 Statistical analysis

Quantified experimental data between different genotypes were analyzed and compared by an unpaired, two tailed Student *t*-test. Data were reported as average ± standard deviation (SD). P-values less than 0.05 were considered as significant.

## ACKNOWLEDGEMENTS

Z.F. and D.L.B. conceived the project. W.X., Z.F., J.S.T designed, constructed, and evaluated the mouse model. A.L., C.C., J.S.T., W.X., Z.F. performed experiments and conducted data curation and analysis. Z.F., D.L.B., W.X. contributed essential reagents/tools and acquired funding support. Z.F., C.C., A.L. contributed to the original writing, D.L.B. and W.X. contributed to the editing of the manuscript.

We thank our colleagues at UVA Genetically Engineered Murine Model Core, Research Histology Core, Advanced Microscopy Facility, and Biorepository Tissue Research Facility for technical support. This work was supported by NIGMS grant GM127690 to Z.F., NCI CCSG 5P30CA044579 to UVA School of Medicine Research Cores, and University of Virginia Harrison Research Grant to C.C.

## FOOTNOTES

KATNIP, Katanin-interacting protein; JS, Joubert syndrome; CSF, cerebrospinal fluids; CILK1, ciliogenesis associated kinase 1; DUF, domain of unknown function; KO, knockout; pH3, phospho-histone H3; ARL13B, ADP-ribosylation factor-like protein 13B; FOP, FGFR1 Oncogene Partner; CRISPR, Clustered Regularly Interspaced Short Palindromic Repeats; Cas12a, CRISPR-associated protein 12a; Cpf1, CRISPR from *Prevotella* and *Francisella*.

## FIGURE LEGEND

Figure 1. : Generation of a *Katnip* knockout mouse model. (A) Schematic illustration of CRISPR/Cas12a-mediated knockout of *Katnip*. (B) Genotyping PCR results showing distinct patterns of the wild type (WT, 267 bp) and the knockout (KO, 242 bp) bands between three different genotypes. (C) A Western blot analysis of Katnip protein in mouse brain tissues from three different genotypes.

Figure 2. : Homozygous depletion of *Katnip* produces a perinatal lethal hydrocephalic phenotype. (A) Representative images of the whole body and the dorsal view of dissected brains from the WT control and the homozygous null littermate at postnatal day 9 (P9). (B) Shown are relative body length, *n* = 4 mice, P9, average ± SD, two-sided unpaired Student’s *t* test, ***P < 0.001. (C) Hematoxylin and eosin (H&E) staining of coronal mouse brain sections, showing broad expansion of the lateral ventricles in the homozygote brain. Scale bar, 500 µm.

Figure 3. : Katnip deficiency decreases the proliferation of neuroprogenitor cells. (A) Representative fluorescence microscopy images showing phospho-histone H3 (pH3)-positive mitotic cells (red), Ki67-positive proliferating cells (red), and Sox2-positive neuroprogenitor cells (green) in the dentate gyrus of wild type (WT) and homozygous knockout (KO) mice. DAPI stains nuclei (blue). Scale bar, 5 µm. (B) Quantification of pH3+ and Ki67+ rates in neuroprogenitor cells from WT and homozygous KO brains (*n* = 3 mice, P9). Shown are average ± SD, two-sided unpaired Student’s *t* test, *P < 0.05.

Figure 4. : Loss of Katnip increases primary cilia frequency and length in the brain. (A) Representative images of primary cilia from WT control and KO mutant mouse hippocampal sections. Cilia axoneme stained with Arl13B (green), basal body stained with FOP (red), and nuclei stained with DAPI (blue). Scale bar, 5 µm. (B). Quantification of cilia frequency and cilia length in hippocampal tissue sections between three pairs of KO mutants and WT littermates (*n* = 3 mice, P9). Shown are average ± SD, two-sided Student’s *t* test, *P<0.05.

Figure S1: *Katnip* null homozygote shows unsteady movements. This movie illustrates that the homozygote mouse (smaller in size) walked unsteadily around the wild mouse, showing signs for loss of balance and coordination in body movement.

## REFERENCES

1. Tran AM, Jnah AJ, De Castro Pretelt MJ. Genetics Review: Joubert Syndrome. Neonatal Netw. Jun 1 2025;44(3):159–166. 10.1891/NN-2024-0052.

2. Glass IA, Dempsey JC, Parisi M, Doherty D. Joubert Syndrome. In: Adam MP, Bick S, Mirzaa GM, Pagon RA, Wallace SE, Amemiya A, eds. GeneReviews((R)). Seattle (WA): 1993.

3. Marshall WF. Basal bodies platforms for building cilia. Curr Top Dev Biol. 2008;85:1–22. 10.1016/S0070-2153(08)00801-6.

4. Mill P, Christensen ST, Pedersen LB. Primary cilia as dynamic and diverse signalling hubs in development and disease. Nat Rev Genet. Jul 2023;24(7):421–441. 10.1038/s41576-023-00587-9.

5. Plotnikova OV, Pugacheva EN, Golemis EA. Primary cilia and the cell cycle. Methods Cell Biol. 2009;94:137–60. 10.1016/S0091-679X(08)94007-3.

6. Wang L, Dynlacht BD. The regulation of cilium assembly and disassembly in development and disease. Development. Sep 17 2018;145(18). 10.1242/dev.151407.

7. Kasahara K, Inagaki M. Primary ciliary signaling: links with the cell cycle. Trends Cell Biol. Dec 2021;31(12):954–964. 10.1016/j.tcb.2021.07.009.

8. Aksu Uzunhan T, Erturk B, Aydin K, et al. Clinical and genetic spectrum from a prototype of ciliopathy: Joubert syndrome. Clin Neurol Neurosurg. Jan 2023;224:107560. 10.1016/j.clineuro.2022.107560.

9. Roosing S, Rosti RO, Rosti B, et al. Identification of a homozygous nonsense mutation in KIAA0556 in a consanguineous family displaying Joubert syndrome. Hum Genet. Aug 2016;135(8):919–921. 10.1007/s00439-016-1689-z.

10. Sanders AA, de Vrieze E, Alazami AM, et al. KIAA0556 is a novel ciliary basal body component mutated in Joubert syndrome. Genome Biol. Dec 29 2015;16:293. 10.1186/s13059-015-0858-z 10.1186/s13059-015-0858-z [pii].

11. Cauley ES, Hamed A, Mohamed IN, et al. Overlap of polymicrogyria, hydrocephalus, and Joubert syndrome in a family with novel truncating mutations in ADGRG1/GPR56 and KIAA0556. Neurogenetics. May 2019;20(2):91–98. 10.1007/s10048-019-00577-2.

12. Niceta M, Dentici ML, Ciolfi A, et al. Co-occurrence of mutations in KIF7 and KIAA0556 in Joubert syndrome with ocular coloboma, pituitary malformation and growth hormone deficiency: a case report and literature review. BMC Pediatr. Mar 12 2020;20(1):120. 10.1186/s12887-020-2019-0.

13. Turner JS, McCabe EA, Kuang KW, et al. The Scaffold Protein KATNIP Enhances CILK1 Control of Primary Cilia. Mol Cell Biol. 2023;43(9):472–480. 10.1080/10985549.2023.2246870.

14. Carpenter EH, Chu CY, Limerick A, Brautigan DL, Fu Z. Human disease variants of KATNIP fail to support CILK1 activation and control of primary cilia. J Cell Sci. Oct 15 2025;138(20). 10.1242/jcs.264056.

15. Boldt K, van Reeuwijk J, Lu Q, et al. An organelle-specific protein landscape identifies novel diseases and molecular mechanisms. Nat Commun. May 13 2016;7:11491. 10.1038/ncomms11491 ncomms11491 [pii].

16. Gafner M, Haddad L, Gupta R, et al. Hydrocephalus associated with a molar tooth sign: A distinct subtype of Joubert syndrome. Dev Med Child Neurol. Jul 2024;66(7):948–957. 10.1111/dmcn.15845.

17. Gana S, Valente EM. Joubert syndrome and hydrocephalus: Further expanding the phenotypic spectrum of a pleiotropic ciliopathy. Dev Med Child Neurol. Jul 2024;66(7):834–835. 10.1111/dmcn.15887.

18. Wallmeier J, Dallmayer M, Omran H. The role of cilia for hydrocephalus formation. Am J Med Genet C Semin Med Genet. Mar 2022;190(1):47–56. 10.1002/ajmg.c.31972.

19. Wallmeier J, Nielsen KG, Kuehni CE, et al. Motile ciliopathies. Nat Rev Dis Primers. Sep 17 2020;6(1):77. 10.1038/s41572-020-0209-6.

20. Duy PQ, Greenberg ABW, Butler WE, Kahle KT. Rethinking the cilia hypothesis of hydrocephalus. Neurobiol Dis. Dec 2022;175:105913. 10.1016/j.nbd.2022.105913.

21. Carter CS, Vogel TW, Zhang Q, et al. Abnormal development of NG2+PDGFR-alpha+ neural progenitor cells leads to neonatal hydrocephalus in a ciliopathy mouse model. Nat Med. Dec 2012;18(12):1797–804. 10.1038/nm.2996.

